# Visualizing and interpreting single-cell gene expression datasets with Similarity Weighted Nonnegative Embedding

**DOI:** 10.1101/276261

**Authors:** Yan Wu, Pablo Tamayo, Kun Zhang

## Abstract

High throughput single-cell gene expression profiling has enabled the characterization of novel cell types and developmental trajectories. Visualizing these datasets is crucial to biological interpretation, and the most popular method is t-Stochastic Neighbor embedding (t-SNE), which visualizes local patterns better than other methods, but often distorts global structure, such as distances between clusters. We developed Similarity Weighted Nonnegative Embedding (SWNE), which enhances interpretation of datasets by embedding the genes and factors that separate cell states alongside the cells on the visualization, captures local structure better than t-SNE and existing methods, and maintains fidelity when visualizing global structure. SWNE uses nonnegative matrix factorization to decompose the gene expression matrix into biologically relevant factors, embeds the cells, genes and factors in a 2D visualization, and uses a similarity matrix to smooth the embeddings. We demonstrate SWNE on single cell RNA-seq data from hematopoietic progenitors and human brain cells.

## Introduction

Single cell gene expression profiling has enabled the quantitative analysis of many different cell types and states, such as human brain cell types (Lake et al. 2016; Lake et al. 2017) and cancer cell states (Tirosh et al. 2016; Puram et al. 2017), while also enabling the reconstruction of cell state trajectories during reprogramming and development (Trapnell et al. 2014; Qiu et al. 2017; Setty et al. 2016). Recent advances in droplet based single cell RNA-seq technology (Macosko et al. 2015; Lake et al. 2017) as well as combinatorial indexing techniques (Cao et al. 2017; Rosenberg et al. 2017) have improved throughput to the point where tens of thousands of single cells can be sequenced in a single experiment, creating an influx of large single cell gene expression datasets. Numerous computational methods have been developed for latent factor identification (Buettner et al. 2017), clustering (Wang et al. 2017), cell trajectory reconstruction (Qiu et al. 2017; Setty et al. 2016), and differential expression (Kharchenko et al. 2014). However, visualization of these high dimensional datasets is still critical to their interpretation. The most common visualization method is t-Stochastic Neighbor Embedding (t-SNE), a non-linear visualization method that tries to minimize the Kullback-Leibler (KL) divergence between the probability distribution defined in the high dimensional space and the distribution in the low dimensional space (Maaten & Hinton 2008; van der Maaten 2014).

t-SNE very accurately captures local structure in the data, ensuring cells that are in the same cluster are close together (van der Maaten 2014). This property enables t-SNE to find patterns that other methods, such as Principal Component Analysis (PCA) (Abdi & Williams 2010) and Multidimensional Scaling (MDS) (Kruskal 1964), cannot (Maaten & Hinton 2008). However, t-SNE often fails to accurately capture global structure in the data, such as distances between clusters, possibly due to asymmetry in the KL divergence metric and the necessity of fine-tuning the perplexity hyperparameter (Maaten & Hinton 2008). This can make interpreting higher order features of t-SNE plots difficult. Additionally, existing visualizations lack biological context, such as which genes are expressed in which cell types, often requiring additional plots or tables for interpretation. While some newer methods such as UMAP address the issue of capturing global structure in the data, no methods, to our knowledge, allow for biological information to be embedded onto the visualization (McInnes & Healy 2018).

## Results

**Figure 1:**
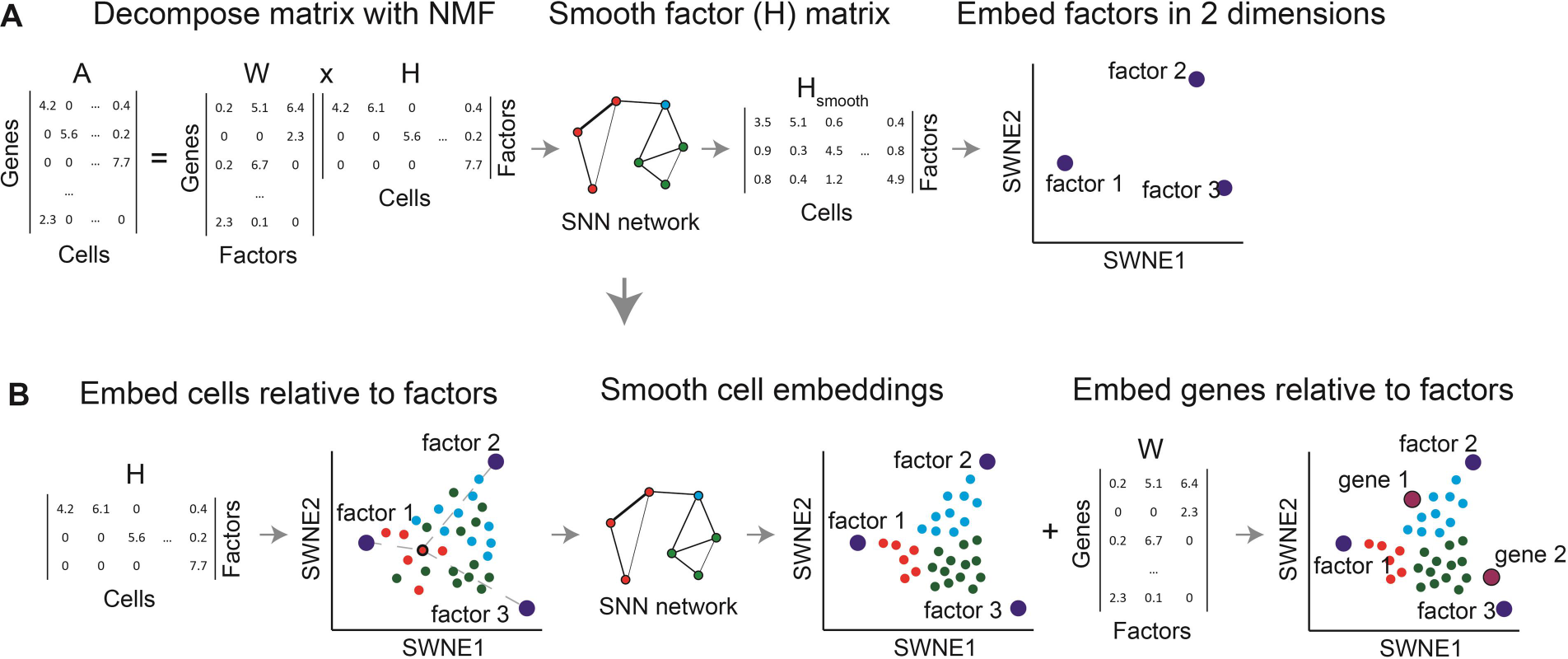
SWNE overview and methodology. **(a)** The gene expression matrix (*A*) is decomposed into a gene loadings matrix (*W*) and a factor matrix (*H*) using NMF, with the number of factors selected via minimizing imputation error (see **Figure S1**). The factor matrix (*H*) is smoothed with the SNN network, and factors (rows of *H*) are embedded in 2 dimensions via Sammon mapping of their pairwise distances. **(b)** Cells are embedded relative to the factors using the cell scores in the *H* matrix, and the cell embeddings are refined using the SNN network. Finally, selected genes are embedded relative to the factors using the gene loadings (*W*).

### SWNE overview and methodology

We developed a method for visualizing high dimensional single cell gene expression datasets, Similarity Weighted Nonnegative Embedding (SWNE), which captures both local and global structure in the data, while enabling the genes and biological factors that separate the cell types and trajectories to be embedded directly onto the visualization. SWNE adapts the Onco-GPS NMF embedding framework (Kim et al. 2017) to decompose the gene expression matrix into latent factors, embeds both factors and cells in two dimensions, and smooths both the cell and factor embeddings by using a similarity matrix to ensure that cells which are close in the high dimensional space are also close in the visualization.

First, SWNE uses Nonnegative Matrix Factorization (NMF) (Lee & Seung 1999; Franc et al. 2005) to create a parts based factor decomposition of the data (**Figure 1a**). The number of factors (*k*) is chosen by randomly selecting 20% of the gene expression matrix to be set to missing, and then finding the factorization that best imputes those missing values, minimizing the mean squared error (Euclidean distance) (**Figure S1a**). With NMF, the gene expression matrix (*A*) is decomposed into: (1) a *genes by factors* matrix (*W*), and (2) a *factors by cells* matrix (*H*) (**Figure 1a**). SWNE then uses the similarity matrix, specifically a Shared Nearest Neighbors (SNN) network (Houle et al. 2010), to smooth the *H* matrix, resulting in a new matrix *H*_*smooth*_. SWNE calculates the pairwise distances between the rows of the *H*_*smooth*_ matrix, and uses Sammon mapping (Sammon 1969) to project the distance matrix onto two dimensions (**Figure 1b**). Next, SWNE embeds cells relative to the factors using the cell scores in the unsmoothed *H* matrix and uses the SNN network to smooth the cell coordinates so that cells which are close in the high dimensional space are close in the visualization (**Figure 1b**). Finally, SWNE embeds genes relative to the factors using the gene loadings in the *W* matrix (**Figure 1b**).

**Figure 2:**
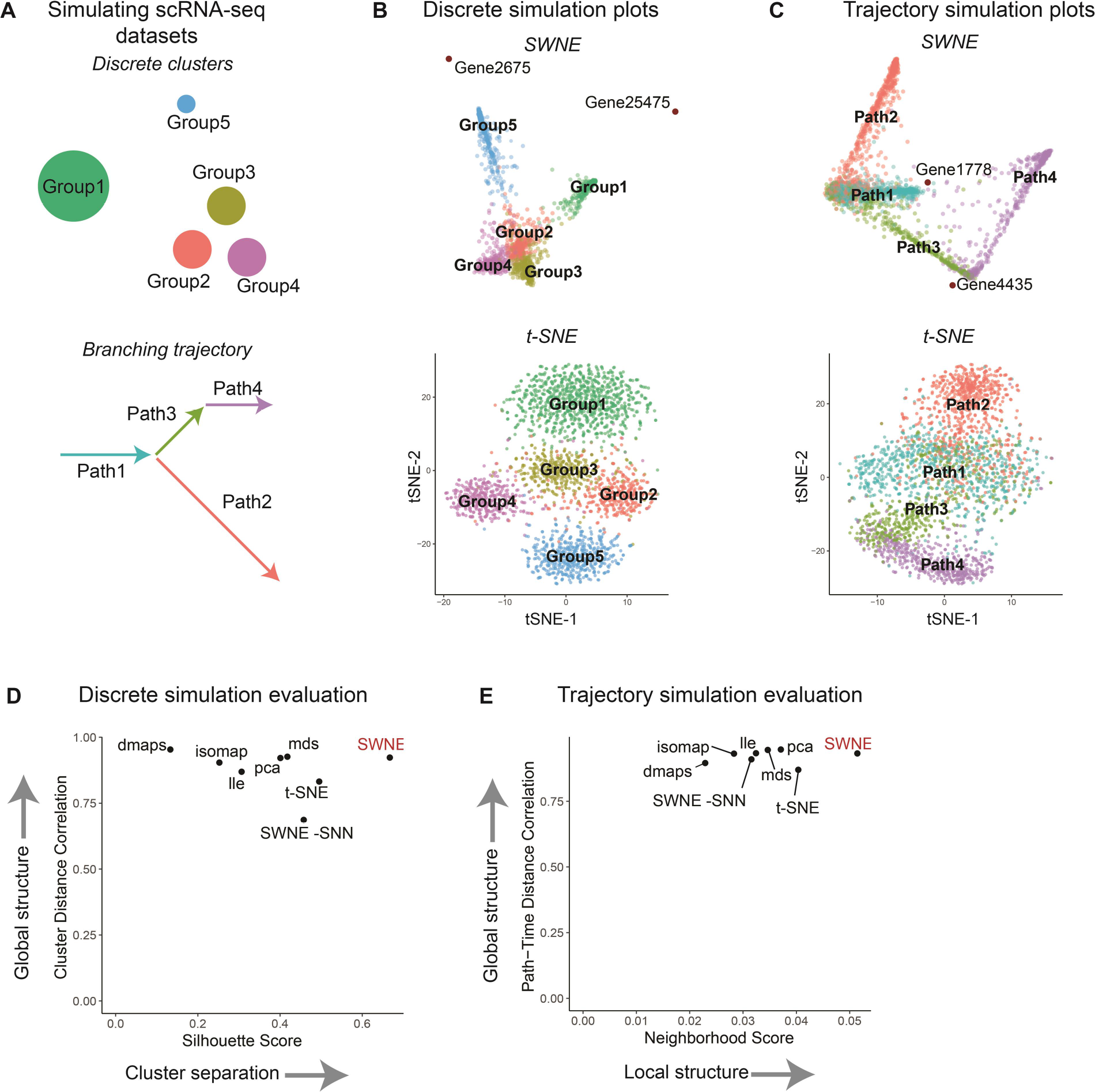
SWNE captures local and global structure in simulated datasets more faithfully than t-SNE and other methods. **(a)** Simulating a discrete dataset with five clusters, and a branching trajectory dataset with four paths. **(b)** SWNE and t-SNE plots of the simulated discrete dataset (see **Figure S2a** for additional plots). **(c)** SWNE and t-SNE plots of the simulated trajectory dataset (see **Figure S2b** for additional plots). **(d)** Quantitative evaluation of SWNE and existing visualization methods on the discrete simulation. Global structure is evaluated by correlating pairwise cluster distances in the embedding with distances in the original gene expression space. Cluster separation is evaluated with the Silhouette score. **(e)** Quantitative evaluation of SWNE and existing visualization methods on the trajectory simulation. Global structure is evaluated by dividing each path up into time steps, and correlating pairwise path-time-step distances in the embedding with distances in the original gene expression space. Local structure is evaluated by taking the Jaccard similarity of the nearest neighbors in the embeddings with the true nearest neighbors.

### SWNE captures local and global structure in simulated datasets more faithfully than t-SNE

To benchmark SWNE against PCA, t-SNE, and other visualization methods, we used the Splatter single-cell RNA-seq simulation method (Zappia et al. 2017) to generate two synthetic datasets. We generated a 2700 cell dataset with five discrete groups, where Groups 2 - 4 were relatively close and Groups 1 & 5 were further apart (**Figure 2a**). We also generated a simulated branching trajectory dataset with 2730 cells and four different paths, where Path 1 branches into Paths 2 & 3, and Path 4 continues from Path 3 (**Figure 2a**). For the discrete simulation, the t-SNE plot qualitatively distorts the cluster distances, making Groups 1 & 5 closer than they should be to Groups 2 – 4 (**Figure 2b**). The SWNE plot more accurately shows that Groups 1 & 5 are far from each other and Groups 2 – 4 (**Figure 2b**). PCA, LLE, and MDS do a better job of accurately visualizing these cluster distances, but have trouble separating Groups 2 – 4 (**Figure S2a**). For the branching trajectory simulation, the t-SNE plot incorrectly expands the background variance of the paths, while the SWNE plot does a much better job of capturing the important axes of variance, resulting in more clearly defined paths (**Figure 2c**). PCA, LLE, and MDS again do a better job of capturing the trajectory-like structure of the data, but still inaccurately expand the background variance more than SWNE (**Figure S2b**).

To quantitatively benchmark the visualizations, we developed metrics to quantify how well each embedding captures both the global and local structure of the original dataset. For the discrete simulation, we calculated the pairwise distances between the group centroids in the original gene expression space, and then correlated those distances with the pairwise distances in the 2D embedding space to evaluate the embeddings’ ability to capture global structure (**Figure 2d**). To evaluate local structure, we calculated the average Silhouette score (Rousseeuw 1987), a measure of how well the groups are separated, for each embedding (**Figure 2d**). SWNE outperforms t-SNE and performs nearly as well as PCA, MDS, and Diffusion Maps in maintaining global structure (**Figure 2d**). SWNE also clearly outperforms every other method, including t-SNE, in cluster separation (**Figure 2d**).

For the trajectory simulation, since we know the simulated pseudotime for each cell, we divide each path into groups of cells that are temporally close **(Methods)**. We then evaluate global structure by calculating pairwise distances between each path-time-group in the original gene expression space and the 2D embedding space, and then correlating those distances (**Figure 2e**). We can evaluate local structure by constructing a ground truth neighbor network by connecting cells from adjacent pseudotimes, and then computing the Jaccard distance between each cell’s ground truth neighborhood matches and its 2D embedding neighborhood **(Methods, Figure 2e**). SWNE outperforms t-SNE in capturing global structure, and performs about as well as PCA, MDS, and LLE (**Figure 2e**). For capturing neighborhood structure, SWNE again outperforms every other embedding, including t-SNE (**Figure 2e**). Finally, both the qualitative and quantitative benchmarks show that SNN smoothing of the cell and factor embeddings is critical to SWNE’s performance, especially for capturing local structure in the data (**Figure 2b, 2c**, **Figure S2a, S2b**).

We used these quantitative metrics to assess how changing the number of factors affects performance for both the trajectory and discrete simulated datasets. The quantitative performance of SWNE is fairly robust across the number of factors used, although there is more of a penalty for using too few factors than too many (**Figure S1b, S1c**). Visually, using too few factors again results in sub-optimal cluster separation, while using too many does not seem to affect performance (**Figure S1d, S1e**). Additionally, we performed a runtime analysis of SWNE on simulated datasets, showing that SWNE scales linearly with the number of cells (**Figure S2c**). SWNE seems to scale in polynomial time with the number of genes, suggesting that feature selection is crucial to reducing runtimes (**Figure S2d**). Using the top 3000 overdispersed genes, the entire SWNE workflow takes about 17 minutes to run on a 50,000 cell dataset (**Figure S2c**).

**Figure 3:**
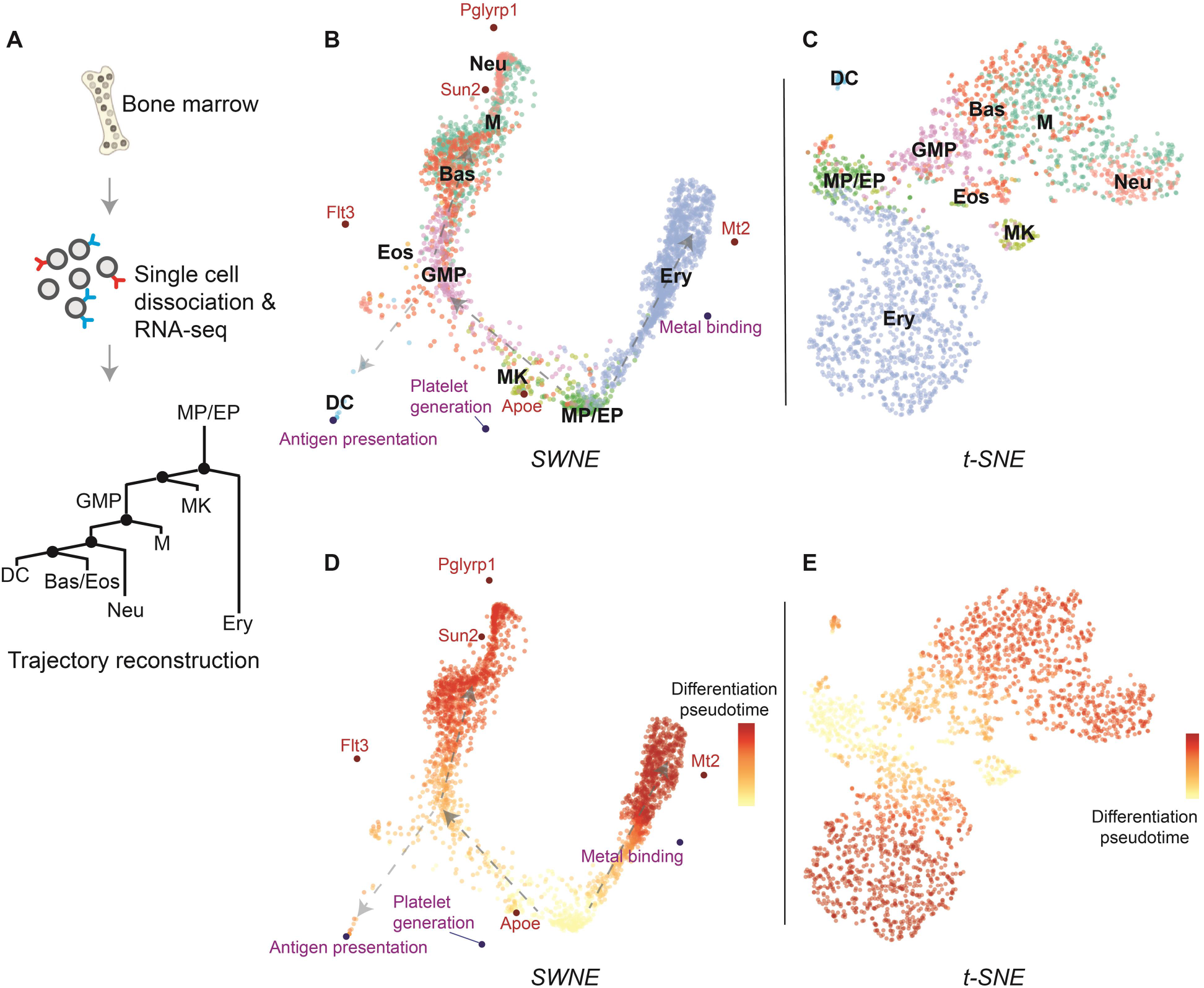
Illuminating the branching structure of hematopoiesis. **(a)** Paul et al sorted single hematopoietic cells from bone marrow, sequenced them using scRNA-seq, and identified the relevant cell types. The hematopoiesis trajectories were reconstructed using Monocle2, and the cells were ordered according to their differentiation pseudotime. **(b)** SWNE plot of hematopoiesis dataset, with selected genes and biological factors displayed (see **Figure S3a-b** for gene and factor annotations). **(c)** t-SNE plot of hematopoiesis dataset. **(d)** SWNE plot of hematopoiesis dataset, with developmental pseudotime calculated from Monocle2 overlaid onto the plot. **(e)** t-SNE plot of hematopoiesis dataset, with developmental pseudotime overlaid onto the plot.

### Illuminating the branching structure of hematopoiesis

We then applied SWNE to analyze the single cell gene expression profiles of hematopoietic cells at various stages of differentiation (Paul et al. 2015) (**Figure 3a**). Briefly, single cells were sorted from bone marrow and their mRNA was sequenced with scRNA-seq (Paul et al. 2015) (**Figure 3a**). The differentiation trajectories of these cells were reconstructed using Monocle2 (Qiu et al. 2017), a method built to identify branching trajectories and order cells according to their differentiation status, or “pseudotime” (**Figure 3a**). The branched differentiation trajectories are shown in the tree in **Figure 3a**, starting from the monocyte and erythrocyte progenitors (MP/EP) and either moving to the erythrocyte (Ery) branch on the left, or the various monocyte cell types on the right (Qiu et al. 2017). We selected the number of factors using our imputation-based model selection method (Figure S1f, **Methods**).

Qualitatively, the SWNE plot (**Figure 3b**) separates the different cell types at least as well as the t-SNE plot (**Figure 3c**). However, the SWNE plot does a much better job of capturing the two dominant branches: the erythrocyte differentiation branch and the monocyte differentiation branch, and shows that those two branches are the primary axes of variation in this dataset (**Figure 3b**). While the t-SNE plot captures the correct orientation of the cell types, it disproportionately expands many of them, obfuscating the branch-like structure of the data (**Figure 3c**). Neither SWNE nor t-SNE accurately orient the different monocyte cell types in the monocyte branch, most likely because the variance is dominated by the erythrocyte - monocyte split, and the extent of differentiation. We also used Monocle2 to calculate differentiation pseudotime for the dataset, which is a metric that orders cells by how far along the differentiation trajectory they are (Qiu et al. 2017). We then overlaid the pseudotime score on the SWNE and t-SNE plots (**Figure 3d, 3e**). Again, we can see that in the SWNE plot, there’s a clear gradient of cells at different stages of differentiation along the two main branches (**Figure 3d**). The gradient in the t-SNE plot is not as visible, most likely because t-SNE obscures the branching structure by expanding the more differentiated cell types (**Figure 3e**).

SWNE provides an intuitive framework to visualize how specific genes and biological factors contribute to the visual separation of cell types or cell trajectories by embedding factors and genes onto the visualization. We used the gene loadings matrix (*W*) to identify the top genes associated with each factor, as well as the top marker genes for each cell type, defined using Seurat (Butler et al. 2018; Satija et al. 2018) (**Methods**). We chose three factors and five genes that we found biologically relevant (**Figure S3a, S3b**). The five genes are: *Apoe, Flt3, Mt2, Sun2*, and *Pglyrp*. The three factors are: Antigen Presentation, Metal Binding, and Platelet Generation, and factor names were determined from the top genes associated with each factor (**Figure S3b**) (**Table S1**). The factors and genes enable a viewer to associate biological processes and genes with the cell types and latent structure shown in the data visualization. For example, dendritic cells (DC) are associated with Antigen Presentation, while erythrocytes (Ery) are associated with Heme metabolism and express *Mt2*, a key metal binding protein (**Figure 3b**). Additionally, the embedded factors and genes allow for interpretation of the overall differentiation process (**Figure 3d**). Undifferentiated progenitors (MP/EP) express *Apoe*, while more differentiated monocytes express *Sun2* and *Pglyrp1* (**Figure 3d**, **Figure S3f**). We validate the location of the *Apoe* embedding by overlaying *Apoe* expression onto the SWNE plot (**Figure S3f**).

**Figure 4:**
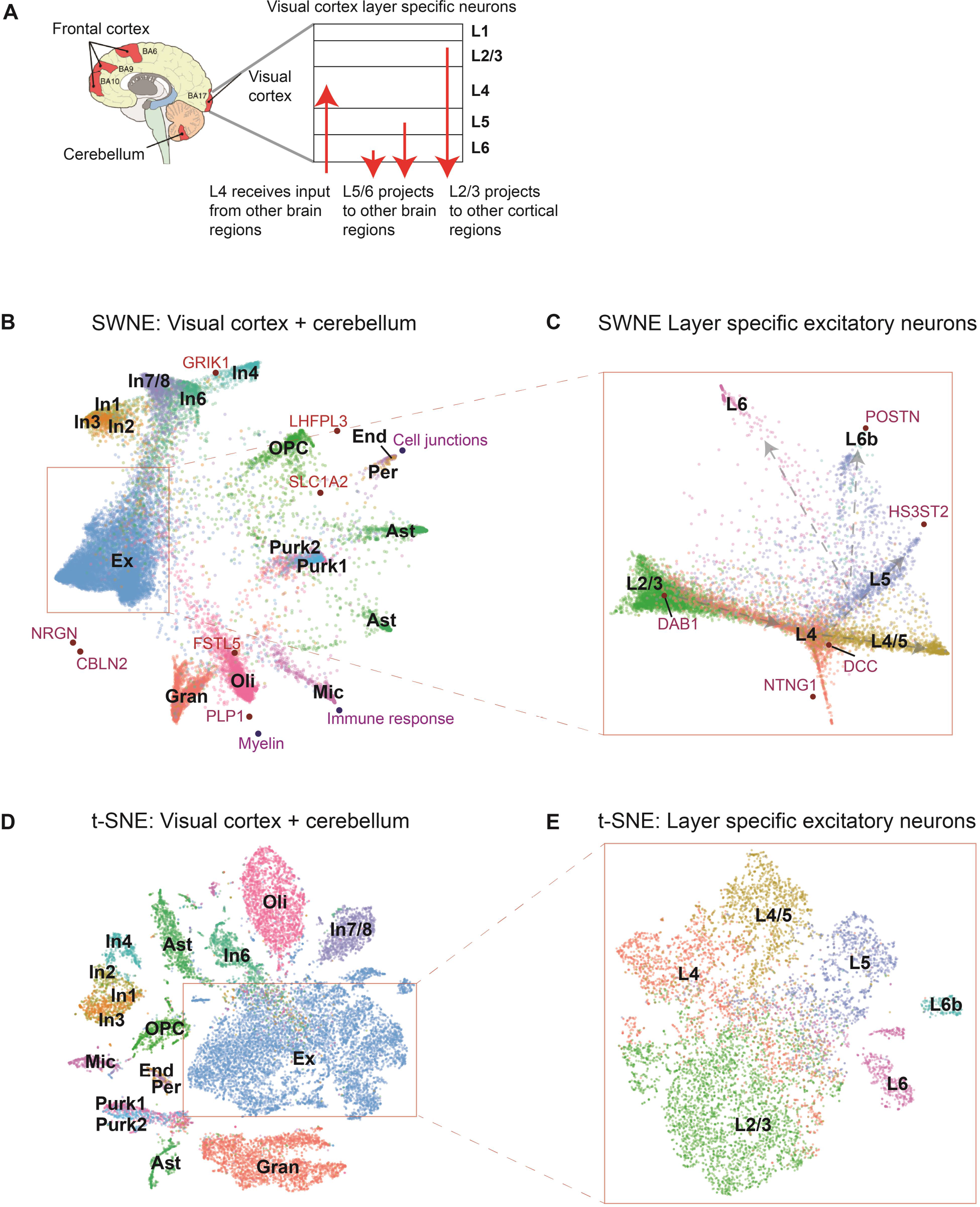
Creating an interpretable map of the human visual cortex and cerebellum. **(a)** Single nuclei were dissociated from the visual cortex and the cerebellum, and sequenced using single nucleus RNA-seq. The inset shows that the excitatory neurons from the visual cortex are grouped into different spatial layers, each of which has different functions. **(b)** SWNE plot of single cells from the visual cortex and cerebellum, with selected genes and factors displayed (see **Figure S3c-d, S3g** for gene and factor annotations). **(c)** Inset: SWNE plot of the excitatory neurons from the visual cortex only, colored by the spatial layer the excitatory neurons belong to (see **Figure S3e, S3h** for gene annotations). **(d)** t-SNE plot of single cells from the visual cortex and cerebellum. **(e)** Inset: t-SNE plot of the excitatory neurons from the visual cortex only, colored by the spatial layer the excitatory neurons belong to.

### Creating an interpretable map of the human visual cortex and cerebellum

We also applied SWNE to a single nucleus RNA-seq human brain dataset (Lake et al. 2017) from the visual cortex (13,232 cells) and the cerebellum (9,921 cells) (**Figure 4a**). Briefly, single nuclei were dissociated from the visual cortex and cerebellum of a single donor and sequenced using single nucleus Drop-seq (Lake et al. 2017). We also applied SWNE to the subset of layer specific excitatory neurons in the visual cortex, where each layer has different functions (Molyneaux et al. 2007; Bernard et al. 2012; Hubel 1995) (**Figure 4a, inset**). To run SWNE, we selected the number of factors using the same missing value imputation method (**Figure S1g, S1h**). We can see that both SWNE (**Figure 4b**) and t-SNE (**Figure 4d**) are able to visually separate the various brain cell types. However, SWNE is able to ensure that related cell types are close in the visualization, specifically that the inhibitory neuron subtypes (In1 - 8) are grouped together at the top of the visualization (**Figure 4b**). t-SNE distorts the distances between inhibitory neurons, visually separating them with Astrocytes (Ast) and the Oligodendrocytes (Oli) (**Figure 4d**).

SWNE again provides an intuitive framework to visualize the contributions of specific genes and factors to the visual separation of cell types. We selected three factors (Myelin, Cell Junctions, and Immune Response) and 8 genes (*PLP1, GRIK1, SLC1A2, LHFPL3, CBLN2, NRGN, GRM1, FSTL5*) to project onto the SWNE plot using the cell type markers and gene loadings (**Figure S3d, S3e**, **Table S1**), adding biological context to the spatial placement of the cell types (**Figure 4b**). We can see that *CBLN2*, a gene known to be expressed in excitatory neuron types (Seigneur & Sudhof 2017), is expressed in the visual cortex excitatory neurons and that *GRIK1*, a key glutamate receptor (Sander 1997), is expressed in inhibitory neurons (**Figure 4b, Figure S3g**). We validate the *CBLN2* embedding by overlaying CBLN2 expression onto the SWNE plot (**Figure S3g**). Additionally, the Myelin biological factor is associated with Oligodendrocytes (Oli), consistent with their function in creating the myelin sheath (Bunge 1968) (**Figure 4b**). The Cell junction biological factor is very close to Endothelial cells (End), reinforcing their functions as the linings of blood vessels (**Figure 4b**).

SWNE has a unique advantage over t-SNE in capturing the local and global structure of the data, exemplified when we zoom into the layer specific excitatory neurons (**Figure 4c, 4e**). SWNE visually separates the different neuronal layers, while also showing that the main axis of variance is along the six cortical layers of the human brain (**Figure 4c**). Each layer seems to branch off of a main trajectory from Layer 4, possibly reflecting Layer 4’s biological function as the only layer to receive input (**Figure 4c**). The t-SNE plot can visually separate the layers, but it is unclear that the main axis of variance is between the different layers (**Figure 4e**). Additionally, we selected five layer specific marker genes (*DAB1, NTNG1, DCC, HS3ST2, POSTN*) to project onto the SWNE plot (**Figure 4c, Figure S3e**). *DAB1*, a signal transducer for Reelin (Trotter et al. 2013), is primarily expressed in Layer 2/3 excitatory neurons, while NTNG1, a protein involved in axon guidance (Lin et al. 2003), is expressed in Layer 4 neurons (**Figure 4c, Figure S3g**). We again validate the position of *DAB1* by overlaying its expression on the SWNE plot (**Figure S3h**).

### Dataset projection

To demonstrate SWNE’s ability to project new data onto existing SWNE embeddings, we used a 3,000 cell PBMC dataset generated by 10X genomics (Zheng et al. 2017), and ran the standard SWNE embedding (**Figure S4a**). We projected a 33,000 cell PBMC dataset with additional cell types onto the training SWNE embedding (**Figure S4b**). The projected embedding obviously cannot distinguish between all the PBMC subtypes since they were not present in the training dataset. Nevertheless, SWNE places the new projected cell types near similar cell types in the training dataset, demonstrating that SWNE embeddings can act as a reference map that new datasets can be projected onto (**Figure S4b**).

## Discussion

### SWNE improves visualization fidelity while adding key biological context

Interpretation and analysis of high dimensional single cell gene expression datasets often involves summarizing the expression patterns of tens of thousands of genes in two dimensions, creating a map that shows viewers properties of the data such as the number of cell states or trajectories, and how distinct cell states are from each other. However while t-SNE, the most popular visualization method, can visualize subtle local patterns of expression that other methods cannot, it often distorts global properties of the dataset such as cluster distances and sizes. This is especially apparent in t-SNE’s inability to visualize time series developmental datasets, as t-SNE tends to exaggerate the size of cell types instead of visualizing the axes of differentiation. Additionally, t-SNE and existing methods only display cells, forcing important biological context, such as the cell type marker genes, to be conveyed in separate plots. Here, we integrate NMF with a Nearest Neighbors smoothing method to create SWNE, a visualization method that preserves global properties of the data and improves upon t-SNE’s ability to capture local properties, while enabling key genes and biological factors to be embedded onto the visualization alongside the cells.

One of SWNE’s key advantages is that the factor embedding framework allows for embedding of genes and cells on the same visualization. The nonnegative factors act as a skeleton for the data, as both cells and genes are embedded relative to these factors. The closer a group of cells is to a gene or a factor on the visualization, the more of that gene or factor the cells express (**Figure S3c, S3g, S3h**). If one thinks of the visualization as a map, these embedded genes and factors act as landmarks, adding key biological context to features of the visualization. Embedding genes and factors also streamlines the presentation of the data, eliminating the need for separate plots of marker genes or genesets. Another key factor in SWNE’s performance is the Shared Nearest Neighbors (SNN) network weighting. Without SNN weighting, the quantitative and qualitative performance of SWNE drops significantly (**Figure 2d, 2e, S2a, S2b**). We believe SNN weighting reduces the effect of biological or technical noise, collapsing the data onto the biologically relevant components of heterogeneity. Surprisingly, this ability to minimize noise enables SWNE to capture local structure in the data even better than t-SNE (**Figure 2d, 2e**). This ability to capture local structure enables SWNE to be particularly effective at illuminating the branch-like structure in developmental trajectory datasets (**Figure 2c, 3b, 3d**).

### Robustness and simplicity of SWNE

One issue with t-SNE is the need to fine-tune the perplexity hyper-parameter, without a cross-validation method to find the optimal perplexity. SWNE also has a critical parameter: the number of factors (*k*) used for the decomposition (**Figure S1a**). However, we include a method for selecting *k*, suggested by the author of the NNLM (Lin & Paul C Boutros 2016) package, which uses NMF to impute missing values in the gene expression matrix (*A*) and selects the *k* that minimizes the imputation error (Figure S1a, **Methods)**. Additionally, SWNE performance is fairly robust across a wide range of *k*. While using too few factors affects SWNE’s ability to separate closely related clusters or trajectories, using too many factors seems to have minimal penalty (**Figure S1b, S1c, S1d, S1e**).

A final factor working in SWNE’s favor is that that the underlying methodology is fairly simple, especially compared to methods such as UMAP and t-SNE (McInnes & Healy 2018; Maaten & Hinton 2008). The Onco-GPS based embedding, and subsequent similarity matrix weighting is very transparent, allowing users to understand how the visualization is being produced. For many users, methods such as UMAP and t-SNE can become a black box, which can result in the incorrect usage of key hyperparameters, such as t-SNE perplexity, and over-interpretation of potential computational artifacts. The simplicity of SWNE also makes it possible to project additional data onto an existing SWNE plot, which is difficult to do with non-linear methods like t-SNE and UMAP (**Methods**, Figure S4a, S4b).

### SWNE limitations and future work

One current limitation of SWNE is the relatively slow runtime when using the whole transcriptome. While we recommend using feature selection which greatly reduces runtime, there are cases when using all genes is necessary. The main computational bottleneck is the NMF decomposition, so future work could focus on improving NMF speed, or substituting NMF with a faster matrix decomposition method such as f-scLVM and pagoda/pagoda2 (Buettner et al. 2017; Fan et al. 2016). Both these methods can also use pre-annotated gene sets to guide the factor decomposition, making interpretation of the factor even easier. A second limitation is that the SNN weighting occurs sequentially after embedding the cells, factors, and genes. This causes the genes and factors to sometimes be further from cell clusters than they should be, although they are still generally closest to the most relevant cell cluster. Future work could involve developing a more elegant method that allows factor embeddings to shift relative to the cell embeddings. This would most likely involve some sort of expectation maximization framework, where the method alternates between optimizing the cell embedding coordinates and the factor embedding coordinates until convergence.

Overall, we developed a visualization method, SWNE, which captures both the local and global structure of the data, and enables relevant biological factors and genes to be embedded directly onto the visualization. Capturing global structure enables SWNE to address issues of distortion that occurs with t-SNE, creating a more accurate map of the data. Capturing local structure with the SNN network smoothing enables SWNE to accurately visualize the key axes of variation. This enables SWNE to illuminate differentiation trajectories and layer-specific neuron structure that is not apparent in other visualizations, such as t-SNE. Finally, embedding key marker genes and relevant biological factors adds important biological context to the SWNE visualization. As single cell gene expression datasets increase in size and scope, we believe that SWNE’s ability to create an accurate, context-rich map of the datasets will be critical to biological interpretation.

## Author Contributions

Y.W., P.T., and K.Z. developed the conceptual ideas and designed the study. Y.W. implemented all computational methods. Y.W., P.T., and K.Z. wrote the manuscript.

## Acknowledgements

We would like to acknowledge Dr. Prashant Mali for his feedback and advice, and Dinh Diep for her technical feedback. Additionally, we would like to acknowledge the Zhang Lab, the Tamayo Lab, and the Mali Lab for their help and support.

Funded in part by NIH grants R01HG009285 (KZ & PT), U01CA217885 (PT), and P30 CA023100 (PT), U01MH098977 (YW & KZ), R01HL123755 (YW & KZ)

## Conflicts of Interest

The authors declare no conflicts of interest.

## Supplementary Figures

**Figure S1:**
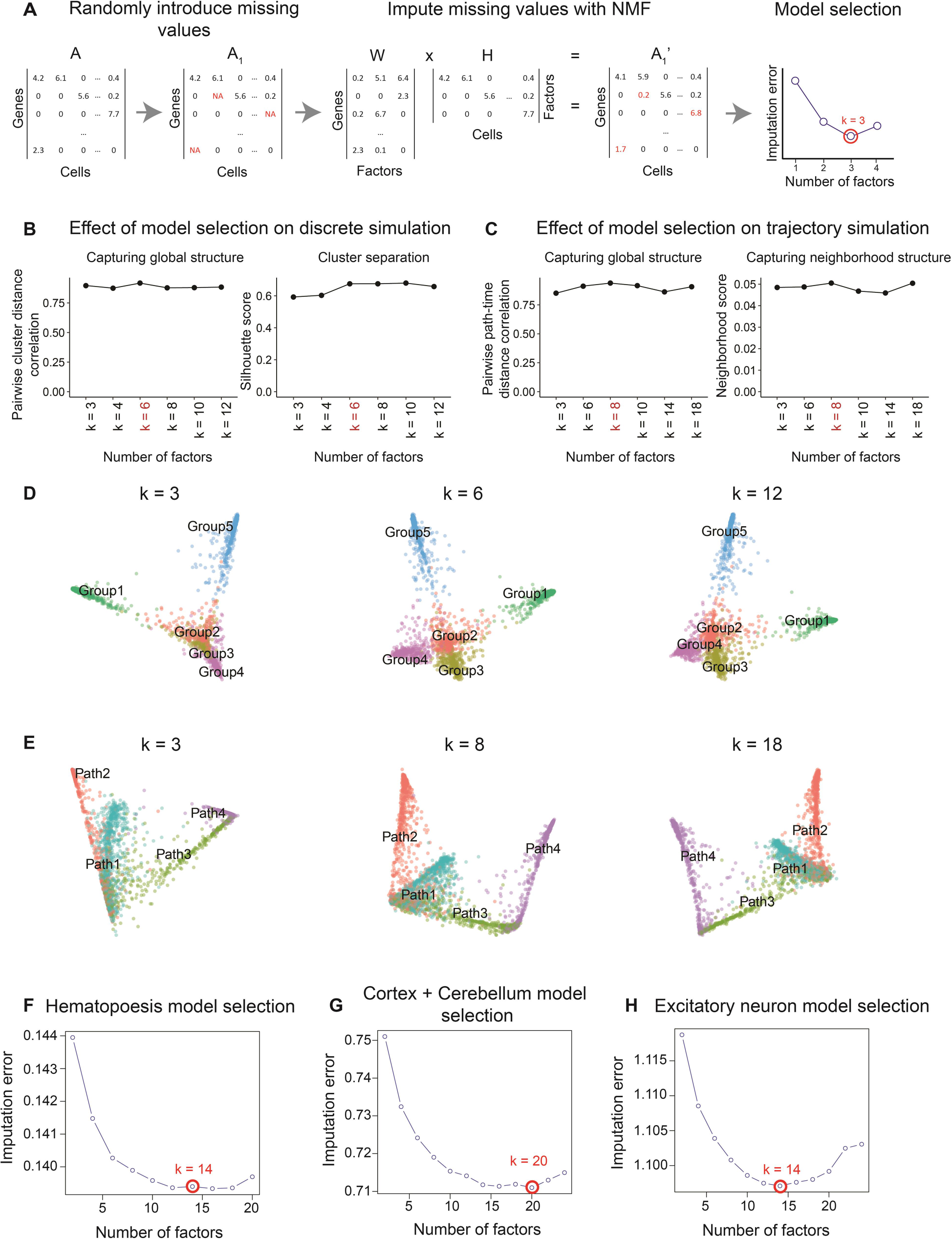
SWNE model selection; related to Figures 1 – 4. **(a)** A subset of the gene expression matrix is set to missing, and NMF is run across a range of factors, and the missing values are imputed. We then plot the imputation error vs number of factors (*k*), and select the *k* values close to the minimum imputation error to create SWNE visualizations with. The *k* that gives the best visualization is selected (**Figure 1**). **(b)** Quantitative evaluation of SWNE performance across a range of *k* for the discrete simulation (**Figure 2**). **(c)** Quantitative evaluation of SWNE performance across a range of *k* for the trajectory simulation (**Figure 2**). **(d)** SWNE visualizations of the discrete simulation across a range of *k* (**Figure 2**). **(e)** SWNE visualizations of the trajectory simulation across a range of *k* (**Figure 2**). **(f)** Imputation error versus *k* for the hematopoiesis dataset (**Figure 3) (g)** Imputation error versus *k* for the cortex & cerebellum dataset (**Figure 3**). **(h)** Imputation error versus *k* for the layer specific excitatory neurons (**Figure 3**).

**Figure S2:**
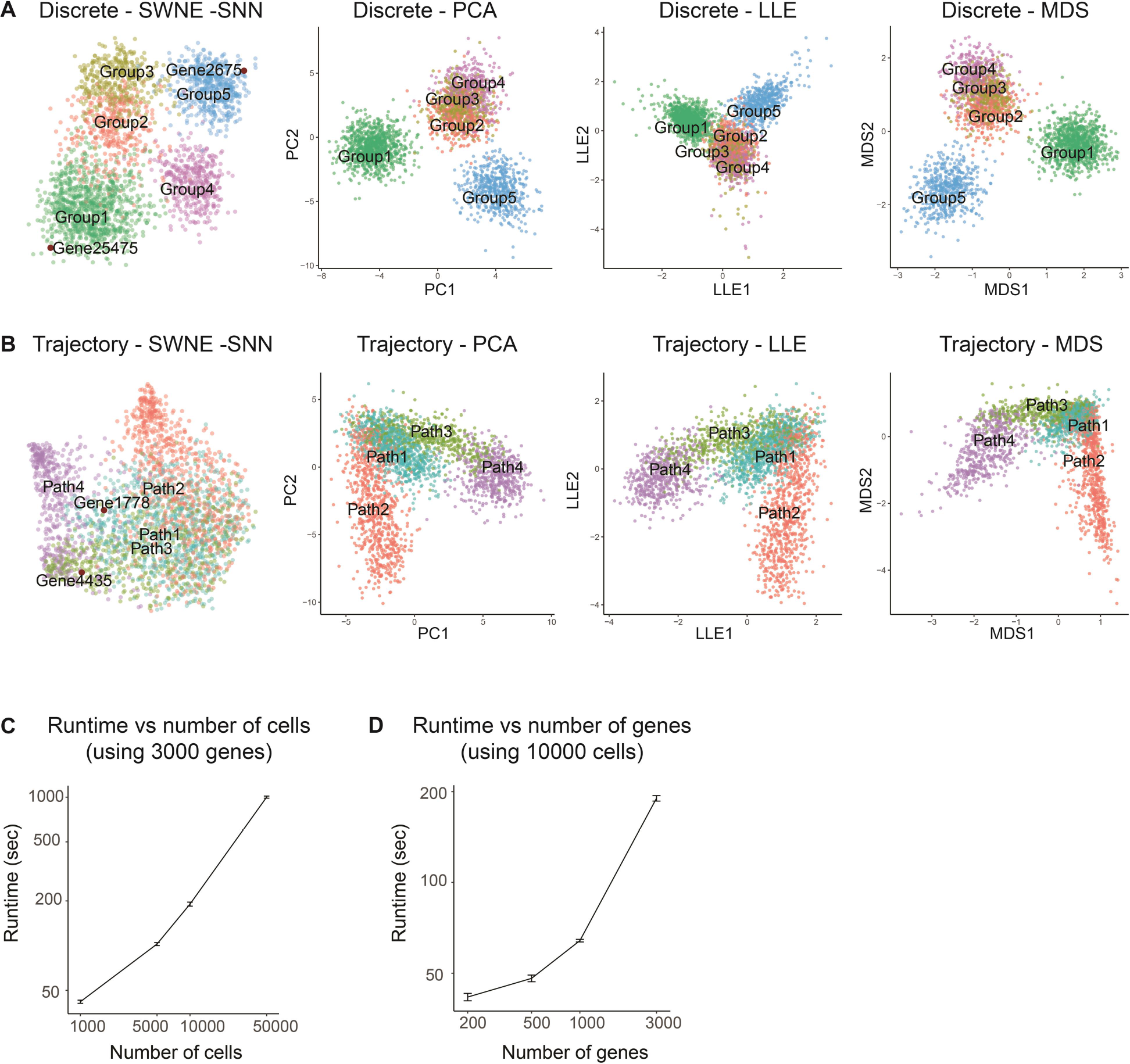
Additional visualizations of the simulated datasets and simulated runtime analysis; related to Figure 2. **(a)** Additional visualizations for the discrete simulation: SWNE without SNN weighting, PCA, locally linear embedding (LLE), multidimensional scaling (MDS). **(b)** Additional visualizations for the trajectory simulation: SWNE without SNN weighting, PCA, locally linear embedding (LLE), multidimensional scaling (MDS). **(c)** SWNE runtime on simulated datasets as a function of the number of cells. **(d)** SWNE runtime on simulated datasets as a function of the number of genes.

**Figure S3:**
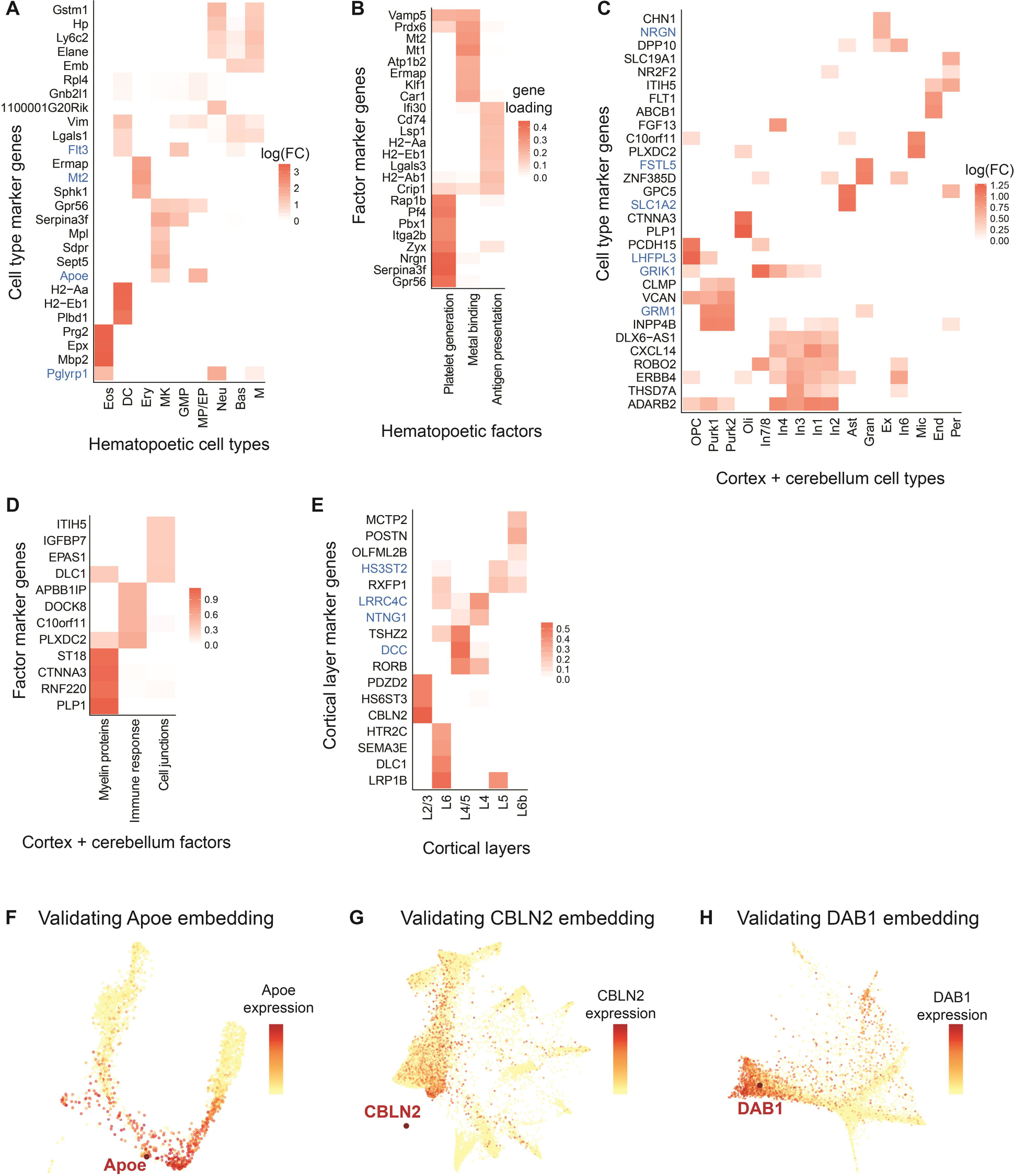
Identifying and validating genes and factors to embed; related to Figure 3 and Figure 4. Genes and factors highlighted in blue are embedded in the corresponding visualization. **(a)** Log fold-change heatmap for the top cell type specific markers in the hematopoiesis dataset (**Figure 2b, Figure 2d**). **(b)** Top gene loadings heatmap for NMF factors in the hematopoiesis dataset (**Figure 2b, Figure 2d**). **(c)** Log fold-change heatmap for the top cell type specific markers in the cortex & cerebellum dataset (**Figure 3b**). **(d)** Top gene loadings heatmap for NMF factors in the cortex & cerebellum dataset (**Figure 3b**). **(e)** Log fold-change heatmap for the top layer specific marker genes in the visual cortex excitatory neuron layers (**Figure 3c**). **(f)** Apoe projected onto the hematopoiesis SWNE plot with Apoe expression overlaid (**Figure 2b, 2d**). **(g)** CBLN2 projected onto the cortex & cerebellum SWNE plot with CBLN2 expression overlaid (**Figure 3b**). **(h)** DAB1 projected onto the layer specific excitatory neuron SWNE plot with DAB1 expression overlaid (**Figure 3c**).

**Figure S4:**
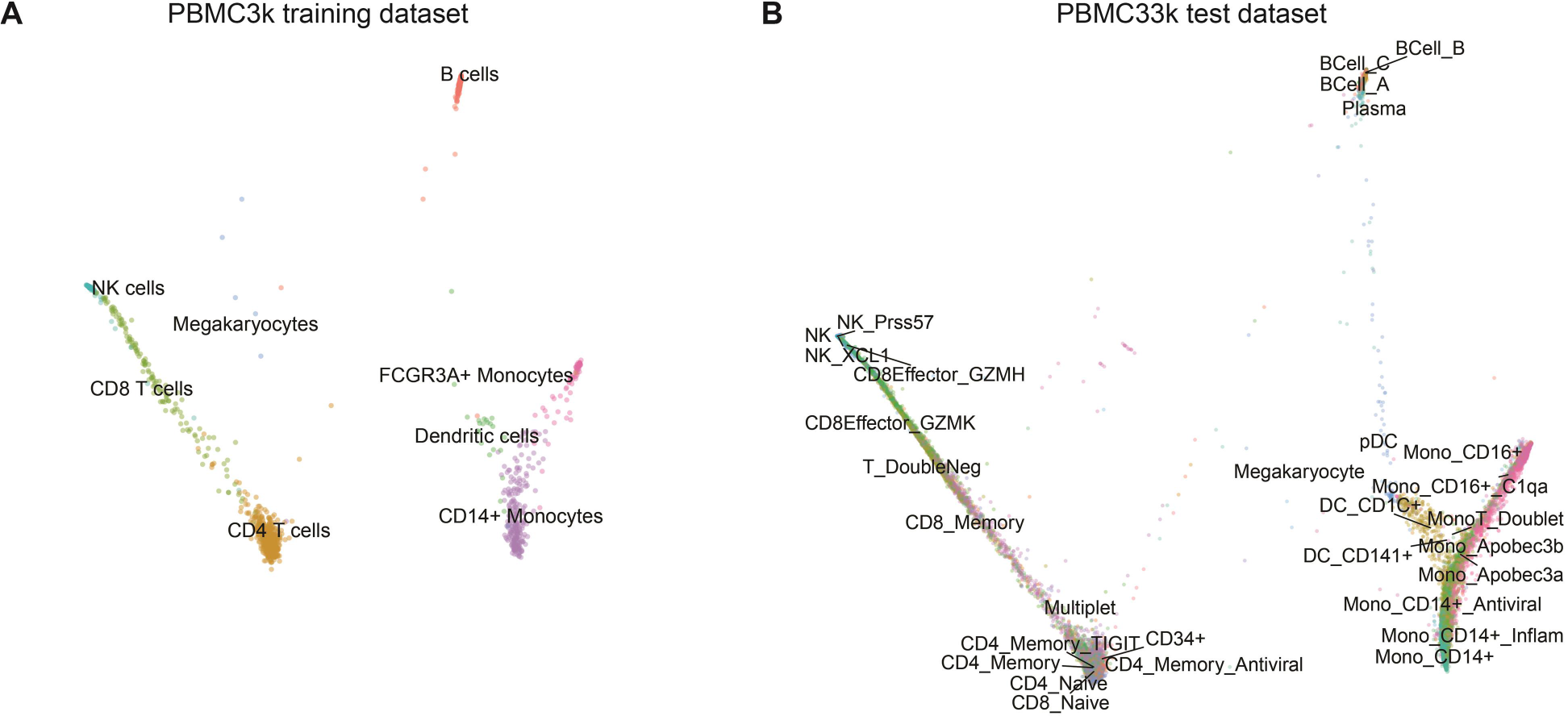
Demonstrating new data projection with PBMCs. **(a)** SWNE was run on a 3,000 PBMC dataset, creating a training embedding. **(b)** SWNE plot of 33,000 PBMCs projected onto the training embedding.

## Method Details

### Normalization, variance adjustment, and scaling

We normalize the gene expression matrix by dividing each column (sample) by the column sum and multiplying by a scaling factor. Batch effects were normalized by a simple model, adapted from pagoda2 (Barkas et al. 2018; Fan et al. 2016), that subtracts any batch specific expression from each gene. We used the variance adjustment method from pagoda (Fan et al. 2016) to adjust the variance of features, an important step when dealing with RNA-seq data. Briefly, a mean-variance relationship for each feature is fit using a generalized additive model (GAM) and each feature is multiplied by a variance scaling factor calculated from the GAM fit. Feature scaling is also performed using either a log-transform, or the Freeman-Tukey transform.

### Feature Selection

We highly recommend doing some sort of selection of overdispersed genes before running SWNE, as the NMF algorithm scales poorly with the number of features (**Figure S2c, S2d**). Both Pagoda2 and Seurat offer dispersion based feature selection methods, and we wrapped with Pagoda2 feature selection method in an SWNE function.

### Nonnegative Matrix Factorization and model selection

We use the NNLM package (Lin & Paul C Boutros 2016) to run the Nonnegative Matrix Factorization (NMF). **Equation 1** shows the NMF decomposition:

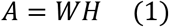

Where *A* is the (features x samples) data matrix, *W* is the (features x factors) feature loading matrix, and *H* is the (factors x samples) low dimensional representation of the data. The NMF initialization method can affect the embedding, and we offer an Independent Component Analysis (ICA) initialization, a Nonnegative-SVD (NNSVD) initialization, and a purely random initialization. ICA initialization works well with most datasets, and is set as the default option. We select the number of factors by setting a random subset of the data as missing, usually around 25% of the matrix, and then use the NMF reconstruction (*W x H*) to impute the missing values across a range of factors. The number of factors, *k*, which minimizes the mean squared error, is typically the optimal number of factors to use.

### Generating the SNN matrix

In order to ensure that samples which are close to each other in the high dimensional space are close in the 2d embedding, we smooth the NMF embeddings with a Shared Nearest-Neighbors (SNN) matrix, calculated using code adapted from the Seurat package (Satija et al. 2018; Butler et al. 2018). Briefly, we calculate the approximate k-nearest neighbors for each sample using the Euclidean distance metric (in the Principal Component space. We then calculate the fraction of shared nearest neighbors between that sample and its neighbors. We can then raise the SNN matrix, denoted here as *S*, to the exponent *β*: *S*′ = *S*^*β*^. If *β* > 1, then the effects of neighbors on the cell embedding coordinates will be decreased, and if *β* < 1, then the effects will be increased. Finally we normalize the SNN matrix so that each row sums up to one.

### Weighted Factor Projection

We adapt the Onco-GPS (Kim et al. 2017) methodology to embed the NMF factors onto a two dimensional visualization. First, we smooth the *H* matrix with the SNN matrix using **Equation 2**:

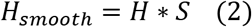

We then calculate the pairwise similarities between the factors (rows of the *H*_*smoo*_ matrix) using either cosine similarity, or mutual information (Kim et al. 2016). The similarity is converted into a distance with **equation 3**:

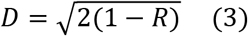

Here, *R* is the pairwise similarity. We use Sammon mapping(Sammon 1969) to project the distance matrix into two dimensions, which represent the x and y coordinates for each factor. The factor coordinates are rescaled to be within the range zero to one.

### Weighted Sample Embedding

Let *F*_*ix*_, *F*_*iy*_ represent the x and y coordinates for factor *i*. To embed the samples, we use the sample loadings from the *unsmoothed H* matrix via **equations 4 & 5**:

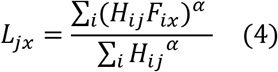

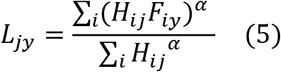

Here, *j* is the sample index and *i* is iterating over the number of factors in the decomposition (number of rows in the *H* matrix). The exponent *α* can be used to increase the “pull” of the NMF components to improve separation between sample clusters, at the cost of distorting the data. Additionally, we can choose to sum over a subset of the top factors by magnitude for a given sample, which can sometimes help reduce noise. We end up with a *2 x N* matrix of sample coordinates, *L*.

To weight the effects of the SNN matrix on the samples, the sample coordinates *L* are smoothed using **equation 6**:

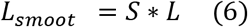

The smoothed sample coordinates (*L*_*smoot*_) are then visualized. While we have found that an SNN matrix works well in improving the local accuracy of the embedding, other similarity matrices, such as those generated by scRNA-seq specific methods like SIMLR, could also work. In general, you should use whichever similarity or distance matrix you used for clustering.

### Embedding features

In addition to embedding factors directly on the SWNE visualization, we can also use the gene loadings matrix (*W*) to embed genes onto the visualization. We simply use the *W* matrix to embed a gene relative to each factor, using the same method we used to embed the cells in the *H* matrix. If a gene has a very high loading for a factor, then it will be very close to that factor in the plot, and far from factors for which the gene has zero loadings.

### Constructing the SNN matrix from different dimensional reductions

The SNN matrix can be constructed from either the original gene expression matrix (*A*), or on some type of dimensional reduction. We have found that constructing the SNN matrix from a PCA reduction tends to work well, especially in datasets where that follow a trajectory or trajectories. We believe this is due to PCA’s ability to capture the axes of maximum variance, while NMF looks for a parts-based representation (Abdi & Williams 2010; Lee & Seung 1999). For datasets where there are discrete cell types, constructing the SNN matrix from the NMF factors is often similar to constructing the SNN matrix from PCA components. Thus, we default to building the SNN matrix from principal components.

### Interpreting NMF components

In order to interpret the low dimensional factors, we look at the gene loadings matrix (*W*). We can find the top genes associated with each factor, in a manner similar to finding marker genes for cell clusters. Since we oftentimes only run the NMF decomposition on a subset of the overdispersed features, we can use a nonnegative linear model to project the all the genes onto the low dimensional factor matrix. One can also run Geneset Enrichment Analysis(Subramanian et al. 2005) on the gene loadings for each factor to find the top genesets associated with that factor.

### Projecting New Data

To project new data onto an existing SWNE embedding, we first have to project the new gene expression matrix onto an existing NMF decomposition, which we can do using a simple nonnegative linear model. The new decomposition looks like **equation 7**:

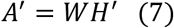

Here, *A*’ is the new gene expression matrix, and *W* is the original gene loadings matrix, which are both known. Thus, we can simply solve for *H*’. The next step is to project the new samples onto the existing SNN matrix. We project the new samples onto the existing principal components, and then for each test sample, we calculate the *k* closest training samples. Since we already have the kNN graph for the training samples, we can calculate, for each test sample, the fraction of Shared Nearest Neighbors between the test sample and every training sample. With the test factor matrix *H*’, and the test SNN matrix, we can run the SWNE embedding as previously described to project the new samples onto the existing SWNE visualization.

### Generating Simulated Datasets

We used the Splatter (Zappia et al. 2017) R package to generate a discrete dataset with five different clusters, estimating parameters from the 3k PBMC dataset published by 10X genomics. We generated five distinct clusters (groups), where Groups 1 and 5 had a differential expressed gene (DEG) probability of 0.3, while Groups 2 - 4 had a DEG probability of 0.15. Group 5 contains 1215 cells, Groups 2 - 4 contain 405 cells each, and Group 1 contains 270 cells. Thus, Groups 1 & 5 should be relatively distant and Groups 2 - 4 should be relatively close. To simulate a branching trajectory dataset, we estimated parameters from the hematopoiesis dataset from Paul et al. We generated four paths, where each path is parameterized by the number of cells in that path and the number of “time-steps”, which essentially controls how long the path is. Path 1 branches into Paths 2 and Paths 3, and Path 3 continues onto Path 4. Paths 1 & 2 contained 819 cells each, and Paths 3 & 4 contained 546 cells each. Path 1 had 100 steps, Path 2 was the “longest” path with 200 steps, and Paths 3 & 4 had 50 steps each. Each cell is assigned to a path, and a time-step. For example, Cell2522 might belong to Path1 and time-step 68.

### Evaluating Embedding Performance

To evaluate how well each embedding maintained the global structure of the discrete simulation, we correlated the pairwise cluster distances in the 2D embedding with the pairwise cluster distances in the original gene expression space. We then calculated the average Silhouette score for each embedding, evaluating how well the visualization separates the clusters. For the trajectory simulation, we divided each path into “chunks” of five time-steps. We correlated the pairwise distances of each “path-time-chunk” in the embedding space, and the original gene expression space to evaluate how well the embeddings maintained the global structure. To evaluate the local structure, we constructed a “ground-truth” neighborhood graph by adding an edge between every cell in each path-time-step, and every cell in each neighboring path-time-step. For example, we would connect all the cells in Path1 at time-step 23, with all the cells in Path1 and time-step 24. We then created a nearest neighbors graph for each embedding, and took the Jaccard distance between each cell’s neighborhood in the embedding and the true neighborhood. We used the average Jaccard distance as our “neighborhood score”.

## Data and Software Availability

The SWNE package is available at https://github.com/yanwu2014/swne. The scripts used for this manuscript are under the Scripts directory. The data needed to recreate the figures can be found here:

- ftp://genome-miner.ucsd.edu/swne_files/hemato_data.tar.gz (Hematopoiesis data)
- ftp://genome-miner.ucsd.edu/swne_files/neuronal_data.tar.gz (Neuronal data)

The raw data for the hematopoietic and neuronal cells can be found at the GEO accessions GSE72857 and GSE97930, respectively. The PBMC dataset can be found at the 10X genomics website: https://support.10xgenomics.com/single-cell-gene-expression/datasets/1.1.0/pbmc3k.

The simulated datasets can be found at:

- ftp://genome-miner.ucsd.edu/swne_files/splatter_simulated_data.tar.gz

## References

Abdi, H. & Williams, L.J., 2010. Principal component analysis. Chemometrics and Intelligent Laboratory Systems, 2, pp.433–459.

Barkas, N. et al., 2018. pagoda2: A package for analyzing and interactively exploring large single-cell RNA-seq datasets. Available at: https://github.com/hms-dbmi/pagoda2.

Bernard, A. et al., 2012. Transcriptional Architecture of the Primate Neocortex. Neuron, 73(6), pp.1083–1099. Available at: http://dx.doi.org/10.1016/j.neuron.2012.03.002.

Blue B. Lake1†, Song Chen1†, Brandon C. Sos1, 4†, Jean Fan2†, Yun Yung3, Gwendolyn E. Kaeser3, 4, Thu E. Duong1, 5, Derek Gao1, Jerold Chun3*, Peter Kharchenko2*, K.Z., 2017. Integrative single-cell analysis by transcriptional and epigenetic states in human adult brain. Nature Publishing Group, (December), pp.1–3. Available at: http://dx.doi.org/10.1038/nbt.4038.

Buettner, F. et al., 2017. f-scLVM: scalable and versatile factor analysis for single-cell RNA-seq. Genome Biology, 18(1), p.212. Available at: https://genomebiology.biomedcentral.com/articles/10.1186/s13059-017-1334-8.

Bunge, R.P., 1968. Glial cells and the central myelin sheath. Physiological Reviews, 48(1), p.197 LP–251. Available at: http://physrev.physiology.org/content/48/1/197.abstract.

Butler, A. et al., 2018. Integrating single-cell transcriptomic data across different conditions, technologies, and species. Nature Biotechnology, (February). Available at: https://www.nature.com/articles/nbt.4096.pdf.

Cao, J. et al., 2017. Comprehensive single cell transcriptional profiling of a multicellular organism by combinatorial indexing. Science, 667(August), pp.1–35.

Fan, J. et al., 2016. Characterizing transcriptional heterogeneity through pathway and gene set overdispersion analysis. Nature Methods, 13(3), pp.241–244.

Franc, V., Hlaváč V. & Navara, M., 2005. Sequential coordinate-wise algorithm for the non-negative least squares problem. Lecture Notes in Computer Science (including subseries Lecture Notes in Artificial Intelligence and Lecture Notes in Bioinformatics), 3691 LNCS(i), pp.407–414.

Houle, M.E. et al., 2010. Can shared-neighbor distances defeat the curse of dimensionality? Lecture Notes in Computer Science (including subseries Lecture Notes in Artificial Intelligence and Lecture Notes in Bioinformatics), 6187 LNCS, pp.482–500.

Hubel, D.H., 1995. Eye, brain, and vision., New York, NY, US: Scientific American Library/Scientific American Books.

Kharchenko, P. V, Silberstein, L. & Scadden, D.T., 2014. Bayesian approach to single-cell differential expression analysis. Nature methods, 11(7), pp.740–2. Available at: http://www.ncbi.nlm.nih.gov/pubmed/24836921.

Kim, J.W. et al., 2016. Characterizing genomic alterations in cancer by complementary functional associations. Nature biotechnology, 34(5), pp.3–5. Available at: http://www.nature.com/doifinder/10.1038/nbt.3527%5Cnhttp://www.ncbi.nlm.nih.gov/pubmed/27088724.

Kim, J.W. et al., 2017. Decomposing Oncogenic Transcriptional Signatures to Generate Maps of Divergent Cellular States. Cell Systems, 5(2), p.105–118.e9. Available at: http://dx.doi.org/10.1016/j.cels.2017.08.002.

Klein, A.M. et al., 2015. Droplet Barcoding for Single-Cell Transcriptomics Applied to Embryonic Stem Cells. Cell, 161(5), pp.1187–1201. Available at: http://linkinghub.elsevier.com/retrieve/pii/S0092867415005000.

Kruskal, J.B., 1964. Multidimensional scaling by optimizing goodness of fit to a nonmetric hypothesis. Psychometrika, 29(1), pp.1–27.

Lake, B. et al., 2016. Neuronal subtypes and diversity revealed by single-nucleus RNA sequencing of the human brain. Science, 357(2013), pp.352–357.

Lee, D.D. & Seung, H.S., 1999. Learning the parts of objects by non-negative matrix factorization. Nature, 401(6755), pp.788–791.

Lin, J.C. et al., 2003. The netrin-G1 ligand NGL-1 promotes the outgrowth of thalamocortical axons. Nature Neuroscience, 6(12), pp.1270–1276.

Lin, X. & Paul C Boutros, 2016. NNLM: Fast and Versatile Non-Negative Matrix Factorization. Available at: https://cran.r-project.org/package=NNLM.

van der Maaten, L., 2014. Accelerating t-sne using tree-based algorithms. The Journal of Machine Learning Research, 15(1), pp.3221–3245. Available at: https://lvdmaaten.github.io/publications/papers/JMLR_2014.pdf%0Ahttp://jmlr.org/papers/v15/vandermaaten14a.html.

Maaten, L. Van Der & Hinton, G., 2008. Visualizing Data using t-SNE. Journal of Machine Learning Research, 9, pp.2579–2605.

Macosko, E.Z. et al., 2015. Highly Parallel Genome-wide Expression Profiling of Individual Cells Using Nanoliter Droplets. Cell, 161(5), pp.1202–1214. Available at: http://linkinghub.elsevier.com/retrieve/pii/S0092867415005498.

McInnes, L. & Healy, J., 2018. UMAP: Uniform Manifold Approximation and Projection for Dimension Reduction. arXiv, pp.1–18. Available at: http://arxiv.org/abs/1802.03426.

Molyneaux, B.J. et al., 2007. Neuronal subtype specification in the cerebral cortex. Nature Reviews Neuroscience, 8(6), pp.427–437.

Paul, F. et al., 2015. Transcriptional Heterogeneity and Lineage Commitment in Myeloid Progenitors. Cell, 163(7), pp.1663–1677. Available at: http://dx.doi.org/10.1016/j.cell.2015.11.013.

Puram, S. V et al., 2017. Single-Cell Transcriptomic Analysis of Primary and Metastatic Tumor Ecosystems in Head and Neck Cancer. Cell, 172, pp.1–14. Available at: https://doi.org/10.1016/j.cell.2017.10.044.

Qiu, X. et al., 2017. Reversed graph embedding resolves complex single-cell trajectories. Nature Methods, 14(10), pp.979–982.

Rosenberg, A.B. et al., 2017. Scaling single cell transcriptomics through split pool barcoding. Bioarxiv.

Rousseeuw, P.J., 1987. Silhouettes: A graphical aid to the interpretation and validation of cluster analysis. Journal of Computational and Applied Mathematics, 20(C), pp.53–65.

Sammon, J.W., 1969. A Nonlinear Mapping for Data Structure Analysis. IEEE Transactions on Computers, C-18(5), pp.401–409.

Sander, T., 1997. Allelic association of juvenile absence epilepsy with a GluR5 kainate receptor gene (GRIK1) polymorphism. American Journal of Medical Genetics - Neuropsychiatric Genetics, 74(4), pp.416–421.

Satija, R., Butler, A. & Hoffman, P., 2018. Seurat: Tools for Single Cell Genomics. Available at: https://cran.r-project.org/package=Seurat.

Seigneur, E. & Sudhof, T.C., 2017. Cerebellins Are Differentially Expressed in Selective Subsets of Neurons Throughout the Brain. J Comp Neurol, 525(15), pp.3286–3311.

Setty, M. et al., 2016. Wishbone identifies bifurcating developmental trajectories from single-cell data. Nature Biotechnology, 34(April), pp.1–14. Available at: http://www.nature.com/doifinder/10.1038/nbt.3569.

Subramanian, A. et al., 2005. Gene set enrichment analysis: a knowledge-based approach for interpreting genome-wide expression profiles. Proceedings of the National Academy of Sciences of the United States of America, 102(43), pp.15545–50. Available at: http://www.ncbi.nlm.nih.gov/pubmed/16199517.

Tirosh, I. et al., 2016. Dissecting the multicellular ecosystem of metastatic melanoma by single-cell RNA-seq. Science, 352(6282), pp.189–196. Available at: http://science.sciencemag.org.gate2.inist.fr/content/352/6282/189.abstract.

Trapnell, C. et al., 2014. The dynamics and regulators of cell fate decisions are revealed by pseudotemporal ordering of single cells. Nature biotechnology, 32(4), pp.381–6. Available at: http://www.ncbi.nlm.nih.gov/pubmed/24658644.

Trotter, J. et al., 2013. Dab1 Is Required for Synaptic Plasticity and Associative Learning. Journal of Neuroscience, 33(39), pp.15652–15668. Available at: http://www.jneurosci.org/cgi/doi/10.1523/JNEUROSCI.2010-13.2013.

Wang, B. et al., 2017. Visualization and analysis of single-cell RNA-seq data by kernel-based similarity learning. Nature Methods, (June 2016), pp.1–6. Available at: http://dx.doi.org/10.1038/nmeth.4207.

Zappia, L., Phipson, B. & Oshlack, A., 2017. Splatter: simulation of single-cell RNA sequencing data. Genome Biology, 18(1), p.174. Available at: http://genomebiology.biomedcentral.com/articles/10.1186/s13059-017-1305-0.

Zheng, G.X.Y. et al., 2017. Massively parallel digital transcriptional profiling of single cells. Nature Communications, 8, pp.1–12. Available at: http://dx.doi.org/10.1038/ncomms14049.

